# Pausing controls branching between productive and non-productive pathways during initial transcription

**DOI:** 10.1101/199307

**Authors:** David Dulin, David L. V. Bauer, Anssi M. Malinen, Jacob J. W. Bakermans, Martin Kaller, Zakia Morichaud, Ivan Petushkov, Martin Depken, Konstantin Brodolin, Andrey Kulbachinskiy, Achillefs N. Kapanidis

## Abstract

Transcription in bacteria is controlled by multiple molecular mechanisms that precisely regulate gene expression. Recently, initial RNA synthesis by the bacterial RNA polymerase (RNAP) has been shown to be interrupted by pauses; however, the pausing determinants and the relationship of pausing with productive and abortive RNA synthesis remain poorly understood. Here, we employed single-molecule FRET and biochemical analysis to disentangle the pausing-related pathways of bacterial initial transcription. We present further evidence that region σ_3.2_ constitutes a barrier after the initial transcribing complex synthesizes a 6-nt RNA (ITC6), halting transcription. We also show that the paused ITC6 state acts as a checkpoint that directs RNAP, in an NTP-dependent manner, to one of three competing pathways: productive transcription, abortive RNA release, or a new unscrunching/scrunching pathway that blocks transcription initiation. Our results show that abortive RNA release and DNA unscrunching are not as tightly coupled as previously thought.

## Introduction

Transcription initiation by DNA-dependent RNA polymerase (RNAP) constitutes the first and often decisive step in gene expression in bacteria. To balance the output of transcription with environmental and cellular needs, an extensive set of molecular mechanisms has evolved to regulate the efficiency and specificity of transcription initiation ^1^. These regulatory mechanisms may either be directly encoded in the transcribed DNA sequence or mediated by protein transcription factors or small-molecule signals. The target of transcription initiation regulators may be the function of RNAP itself, or the accessibility or affinity of promoters for RNAP. Further regulation occurs in the elongation and termination phases of transcription ^2-5^.

To perform promoter-specific transcription initiation, the five-subunit bacterial RNAP core associates with housekeeping σ^70^ initiation factor (or one of the alternative σ factors) to form an RNAP holoenzyme ^6,7^. The RNAP holoenzyme employs sequence-specific interactions between the σ^70^ and the −35 and −10 promoter elements (**Fig. 1A**) to form an initial RNAP-DNA closed complex (RP_C_), and to isomerize to the catalytically-competent RNAP-promoter DNA open complex (RP_O_) ^8,9^ (**Fig. 1B**). During initial RNA synthesis, strong interactions with the DNA hold the RNAP immobile at the promoter, resulting in the build-up of “scrunching” of downstream DNA, a conformational change that increases the size of the DNA bubble ^10-13^. The eventual break-up of RNAP–promoter contacts and the escape to elongation relax the scrunched DNA ^11^. The productive promoter escape pathway competes with abortive initiation, an unproductive pathway wherein the short nascent RNA is thought to prematurely dissociate, resetting the initially transcribing complex (ITC) to RP_O_. The presence of the abortive pathway is firmly established by *in vitro* biochemistry ^14-16^, single-molecule biophysics ^11,17^ and *in vivo* studies ^18^. While conformational strain resulting from the DNA scrunching may promote abortive initiation ^11^, multiple other factors – such as the presence of the σ_3.2_ region (which obstructs the RNA-exit channel; ^19-22^), strong RNAP–promoter interactions ^9,16,23^ and the initially transcribed sequence ^24,25^ – also contribute.

**Fig. 1.**
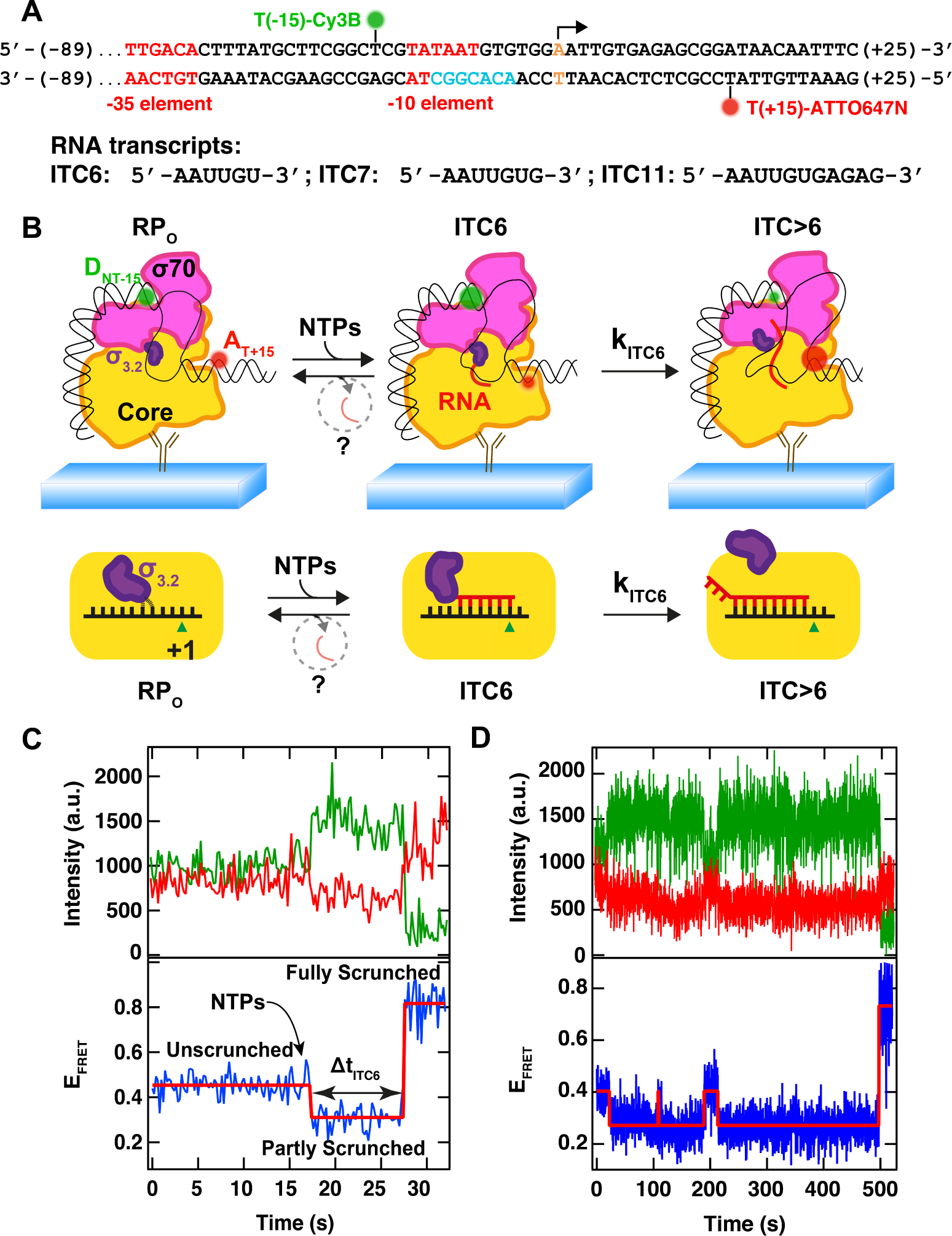
Initial transcription monitored at the single-molecule level. **(A)** Representation of the premelted (turquoise font) WT DNA promoter used in the single-molecule experiments (**Fig. S1A**). The −35 and −10 elements are represented in red. The promoter was donor labeled at −15 position (green sphere) of the non-template DNA strand and acceptor labeled in +15 (red sphere) position of the template DNA strand. An arrow above the base in orange font indicates the +1 position. All the promoters used in the study are described in **Fig. S1A**. **(B)** Schematic of the initial transcription experiment (**Materials and Methods**). Above: using TIRFM-based smFRET, we monitored the E_FRET_ variations of the donor-acceptor pair upon NTP addition. The RNAP fluctuates between RP_O_, ITC6 and ITC>6, to eventually escape the initiation phase towards the elongation phase, or to release the nascent RNA; below: cartoon that magnifies the interactions between the 5’-RNA end and σ3.2 and the position of the 5’-RNA end. **(C)** Fluctuations in the donor (green) and acceptor (red) dyes intensities (above) and the resulting E_FRET_ (below, blue), showing the variation of E_FRET_ from an Unscrunched (US) FRET state, followed by the Partly Scrunched (PS) FRET state upon NTP addition, and ending in the Fully Scrunched (FS) FRET state. Experimental conditions: 200 ms time points (100 ms ALEX, **Materials and Methods**), 500 μM ApA and 80 μM All NTP. **(D)** Similar experiment conducted as described in (C), with the RP failing multiple times before reaching the FS FRET state. Experimental conditions: 200 ms time points, 500 μM ApA and 30 μM All NTP and WT promoter. The red solid lines in the lower panels in (C) and (D) represent the FRET states extracted from a Hidden Markov Modeling (HMM) (**Materials and Methods**).

The step that defines the overall rate of transcription initiation varies between promoters ^9,16,23^. Most σ^70^ promoters in *Escherichia coli* are rate-limited by the stability of RP_C_ or the rate of its isomerization to RP_O_; however, in many cases, the rate-limiting step is attributed to the half-life of RP_O_ or the rate of promoter escape. An extensively studied example of an escape-limited promoter is *lacUV5* ^26^, which is known to produce substantial amounts of abortive products; further, transcriptional pausing has been identified in initially transcribing complexes after the synthesis of 6-nt RNA, at least partly due to the clash of the 5’-RNA end with σ_3.2_ region ^27,28^.

Recent advances in structural characterization of bacterial transcription initiation complexes have created intriguing hypotheses on how specific molecular interactions and conformational changes drive holoenzyme formation, promoter recognition, isomerization to open complex ^29^ and initial RNA synthesis ^12,20,30^. Complementing this fresh structural insight with detailed functional analysis is hampered, however, by the multi-step, asynchronous nature of transcription initiation pathways. Single-molecule techniques, which can provide a direct readout for several steps in the mechanism and resolve co-existing reaction pathways, are well-positioned to overcome the complexity of transcription initiation.

Here, we combined single-molecule and biochemical analysis of initial transcription to explore the mechanistic basis of the pause encountered by ITC6 on *lac* promoter ^27^. We present evidence that the ITC6 pause represents a major control point where the initially transcribing complexes branch to three competing downstream reaction pathways: pause exit by productive transcription; abortive-RNA release; and slow cycling between DNA conformations with different extents of scrunching but without RNA release. The partitioning between these three paths and their kinetics depended on distinct interactions and structural elements. The rate of productive pause exit is synergistically controlled by the initial transcribed sequence and the interaction of the 5’-RNA end with σ_3.2_ region, whereas weak RNAP–promoter interactions favor the entry into the scrunching/unscrunching pathway.

## Results

### High-resolution, real-time observation of initial transcription using single-molecule FRET

To monitor the kinetics of transcription initiation at the single-molecule level, we developed a FRET sensor for real-time imaging of individual RNAP-promoter DNA complexes engaged in nascent RNA synthesis at a *lac* promoter derivative ^27,31^. A similar approach had recently revealed the presence of a strong pause after the synthesis of 6-nt RNA by ITCs ^27,31^. To allow an in-depth biophysical analysis of this rate-limiting ITC6 pause, we modified the original promoter design in two ways (**Fig. 1A** and **Fig. S1A**). First, we extended the upstream region of the promoter DNA fragment from −39 to −89 to enhance RP_O_ formation and provide a more native DNA-length context for RNAP-DNA interactions (Vo et al, 2003; Ross and Gourse, 2005). Second, we moved the acceptor dye from position +20 to +15 to create a labeled promoter DNA fragment that acts as a FRET sensor with clearly separated FRET readouts for three structural states: the open complex RP_O_, the paused complex ITC6, and a pause-cleared complex (ITC11). Since the FRET pair flanks the transcription bubble, the FRET pair reports on the magnitude of DNA scrunching ^10^ during initial RNA synthesis: the point of attachment of the upstream dye (Cy3B) remains fixed when a promoter-anchored RNAP engages in initial RNA synthesis, while the dye (ATTO647N) placed on the DNA downstream of the transcription bubble is pulled towards the active site (**Fig. 1B**). However, each cycle of RNA extension does not decrease the inter-dye distance (or consequently increase the FRET efficiency E_FRET_); the downstream dye at some positions moves farther along from the donor due to the rotation of the downstream dsDNA.

We calibrated the FRET sensor by measuring E_FRET_ for initial transcribing complexes in the presence of different subsets of nucleotides that allowed maximal transcript lengths of 6, 7 or 11 nucleotides, and compared their FRET profiles with RP_O_. These experiments allowed unambiguous assignment of the RP_O_ and ITC11 FRET levels (E_FRET_) as ∼0.5 and ∼0.76, respectively (**Fig. 1C;** see also **Fig. S1**). For clarity, we define these FRET states as “unscrunched” (US) and “fully scrunched” (FS), respectively. Though it is commonly accepted that RNAP escapes *lac* promoters after synthesizing an 11-mer ^9,32^, the high FRET signal after forming a 11-nt long transcript (**Fig. 1C**) is consistent with extended scrunching of the downstream DNA in an ITC (and not an elongation complex). On the other hand, complexes with maximal transcript lengths of 6 and 7 nucleotides yielded E_FRET_ 0.38 and 0.37, respectively. The assignment of this E_FRET_ level to an ITC6 pause state, or “partly scrunched” (PS), was based on our previous observations showing that ITC6 (but not ITC7) accumulates in significant amounts in conditions that allow continuous transcription past the 6–7-mer RNA and that ITCs do not pause again until reaching the position +12 of the DNA template used here ^27^.

Upon addition of NTPs to RP_O_ complexes (either using a NTP subset sufficient to reach the ITC11 complex, or using all NTPs), the FRET signal showed an almost instant transition from the US to the PS state (**Fig. 1C Fig. S1B**), suggesting that the transcription complexes synthesized 6-mer RNA and paused. After the pause, the ITCs split into two main populations: the first population comprised “productive” ITCs that resumed transcription and progressed from the PS to the FS state by synthesizing an 11-mer (**Fig. 1C**). The second population comprised ITCs that returned from the PS to the US state (**Fig. 1D**); notably, such complexes could cycle multiple times (e.g., at ∼100 and ∼200 s in **Fig. 1D**,) between PS and US states until they eventually reached the FS state (e.g., at ∼500 s in **Fig. 1D**).

### Transcribed sequence and 5’-RNA end determine the lifetime of ITC6 pause

Two elements appear to contribute to RNAP pausing at ITC6: *i)* the clash of 5’-RNA end with the σ_3.2_ region (**Fig. 1B**), which blocks the RNA-exit channel of RNAP ^27^, and *ii)* a specific sequence motif (a non-template Y+^6^G+^7^ in the transcribed DNA strand; ^31^) akin to that causing sequence-specific pausing in elongation ^33-35^. We dissected the contributions of these two elements on the ITC6 pause using our smFRET assay.

To explore the steric-clash hypothesis, we modified the 5’-RNA end of the nascent transcript (and thus its interaction with σ_3.2_) by initiating transcription either using ATP or using a synthetic dinucleotide (ApA). The use of ATP as an initiating nucleotide introduces a 5’-triphosphate tail and a net charge of −4 to the 5’-RNA end; in contrast, ApA-primed reactions result in RNAs with no 5’-triphosphate tail. To evaluate the effect of the pause sequence motif on the dynamics of initial transcription, we replaced the sequence T+^6^G+^7^ (on non-template DNA) with G+^6^T+^7^, creating a “ΔP promoter” (**Fig. S1A**) – this substitution has shown to shorten the pause by five-fold in a bulk gel assay and to reduce the total time spent in initial transcription by ∼2-fold in a single-molecule assay ^31^. In all experiments, the initiating ATP or ApA were held at 500 μM, a level significantly above the *K*_M_ of the RP_O_ for initiating nucleotides ^22^; we also varied the concentration of remaining NTPs (1–500 μM).

We first analyzed the effects of the pause elements on the pause duration at ITC6 (Δt_ITC6_) by focusing on the subpopulation of molecules displaying a US→PS→FS scrunching sequence (as in **Fig. 1C)**. The dwell-time distribution for the ITC6 pause was well described by a single exponential (**Fig. 2A**). The pause exit rate (*k*_ITC6_) towards productive synthesis (e.g. formation of ITC11), was extracted using a Maximum Likelihood Estimation (MLE) fitting routine and the errors were evaluated by bootstrapping (for details, see **Materials and Methods**). We observed that in the presence of ApA, the ITC6 pause exit rate was ∼1.5-fold lower for the WT promoter compared to the ΔP promoter (**Fig. 2B**). When we replaced ApA with ATP as the initiating nucleotide and employed the remaining NTPs at above 30 μM, the ITC6 pause exit rate increased from ∼0.07 to ∼0.3 s^-1^ for the ΔP promoter and from ∼0.04 to ∼0.11 s^-1^ for the WT promoter, i.e. the pause exit rate enhancement was more than 2.5-fold (**Fig. 2B**). These experiments demonstrate that the ITC6 pause duration is controlled both by the transcribed sequence and by the 5’-RNA interaction with σ_3.2_.

**Fig. 2.**
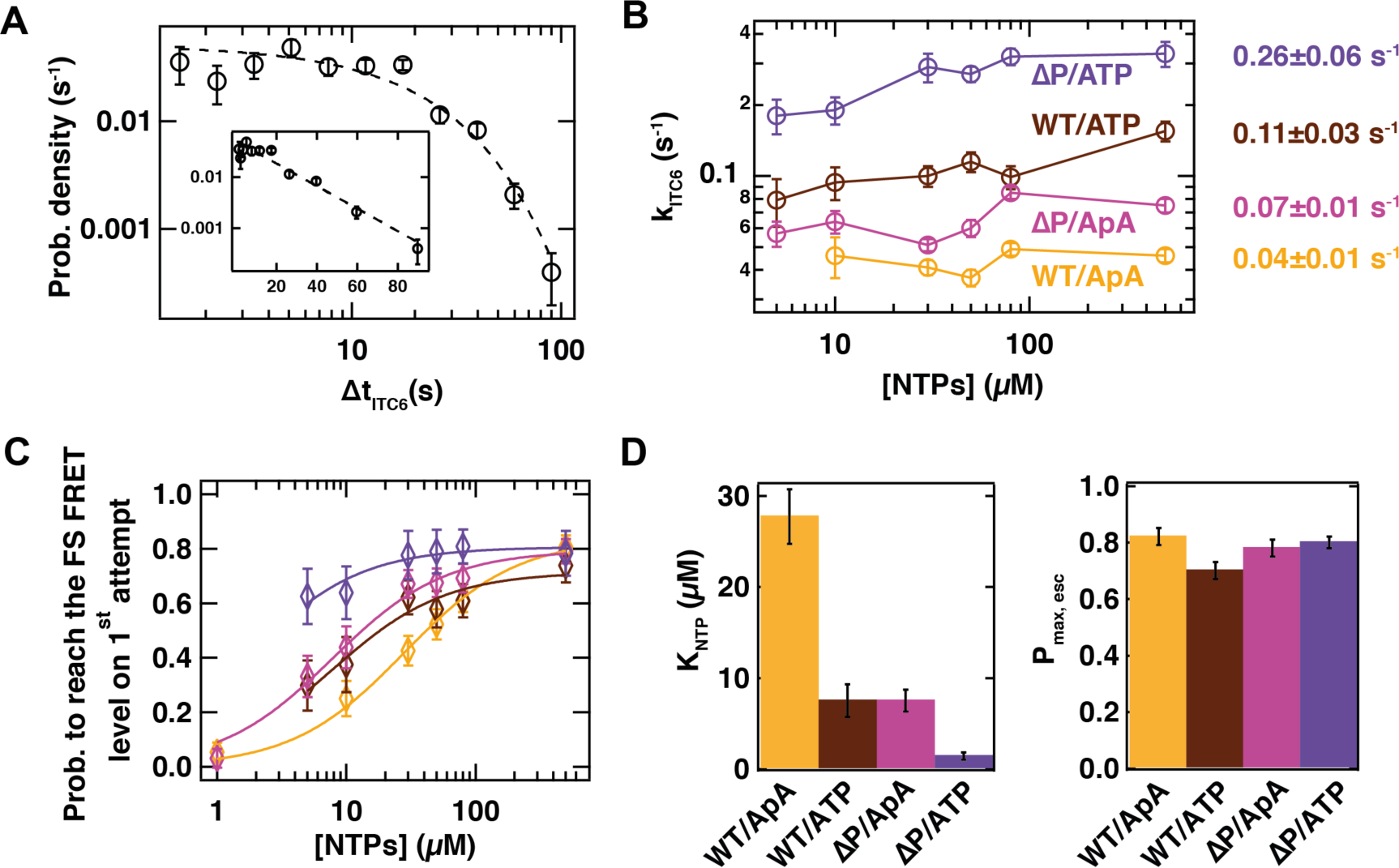
The pause characteristics are a function of the downstream DNA promoter sequence at the +6 position and of the 5’-RNA end nature. **(A)** Probability density distribution for the Δt_ITC6_ for the RP complexes behaving as in **Fig. 1C**. The dashed line is a single-exponential fit from a MLE. Inset: Log-lin representation of the same data. Experimental conditions: 500 μM ApA starting substrate, 80 μM all NTP. **(B)** Δt_ITC6_ lifetime extracted from single exponential MLE fit similar to (A) for different promoter/starting substrate conditions (as indicated in the panel), different NTP conditions, i.e. all NTPs for WT/ApA (yellow) and ITC11 (**Fig. S1A**) for all others, and different NTP concentrations (**Table S1**). In the ATP-initiated reactions, we did not use NTP concentration below 5 μM to prevent potential misincorporations of ATP (used at 500 μM for initiation purposes) ^76^. On the right hand side is indicated the mean ± standard deviation of k_ITC6_ for each promoter/starting substrate condition. **(C)** Probability to reach the fully scrunched (FS) FRET level in a single attempt (**Fig. 1C**). The solid lines are fits to a binding isotherm of the form *p* (*NTP*)= *P*_*max,esc*_× [*NTP*] ([*NTP*]+ *K*_*NTP*_). The error bars are 95% confidence intervals. **(D)** *K*_*NTP*_ and *P* _*max,esc*_×extracted from (C). Error bars are one standard deviation extracted from the fit.

We also noted that the NTP concentration did not influence significantly the ITC6 pause exit rate for the WT promoter with either ApA or ATP as the starting substrate, or for the ΔP promoter with ApA as the starting substrate (**Fig. 2B**). The rate-limiting step in all these cases is thus neither the intrinsic catalytic activity of the transcription complex nor the binding of the incoming NTP substrate. Interestingly, we observed a drop in the ITC6 pause exit rate (from ∼0.3 to ∼0.2 s^-1^) on ATP starting substrate ΔP promoter when the NTP concentration was decreased from 30 to 10 μM. We interpreted these changes as kinetic dependence on two rate-limiting steps for the ΔP + ATP case. The slower, NTP-dependent step, dominant below 10 μM NTP, may be linked to the accessibility of the active site for the incoming NTP, whereas the faster, NTP-independent step, dominant above 30 μM NTP, maybe linked to the structural rearrangement of the interacting σ_3.2_ and the 5’-RNA end.

We next characterized the probability to exit the ITC6 pause on the first attempt (**Fig. 2C**). For this purpose, we counted the probability of ITCs to proceed via the reaction path depicted in **Fig. 1C** (single-scrunch pathway), with or without a detectable ITC6 pause, versus the path in **Fig. 1D** (cyclic scrunching/unscrunching pathway). For the ATP-initiated ΔP promoter (**Fig. 2C**), the probability to exit on the first attempt was high (0.6–0.8) at all substrate NTP concentrations (5–500 μM). On the contrary, the pause-exit probability for the ApA-initiated ΔP promoter, and the ATP- or ApA-initiated WT promoter decreased steeply from 0.8 towards zero at low NTP concentrations (**Fig. 2C**). By fitting the probability *p NTP* to exit the ITC6 pause on the first attempt with a descriptive model similar to a binding isotherm (**Fig. 2C**), we extracted a binding constant *K*_*NTP*_ and a maximal pause-exit probability *P*_*max,esc*_ for each condition (**Fig. 2D**). Overall, the WT promoter had a higher *K*_*NTP*_ compared to ΔP promoter complexes (∼28 vs 8 μM, ApA) while ATP-initiated complexes had a lower *K*_*NTP*_ compared to ApA-initiated ones (∼8 vs ∼28 μM, WT promoter). The probability *P*_*max,esc*_ was relatively constant, with ∼80% of the molecules reaching the FS FRET level on the first attempt at saturating NTP concentration. These results suggest that ITCs can exit a weak ITC6 pause (ΔP promoter) efficiently even at low NTP concentration, while overcoming a strong ITC6 pause (WT promoter + ApA at the 5’-RNA end) requires higher NTP concentration (**Fig. 2C**).

Interestingly, we observed that 3–20% of the ITCs did not display a pause in the PS state (plain bars, **Fig. S2B**), but rather a direct transition from US to FS. This indicates two possible origins for the apparent absence of pausing: the presence of a non-pausing population of ITCs, and inadequate temporal resolution to capture the fastest US→PS→FS transitions. We thus calculated the fraction of the pausing ITCs that we cannot technically detect (by integrating the pause-exit probability distribution from 0 to our detection limit), and subtracted this fraction from the total non-pausing ITC population (plain bars, **Fig. S2B**). The corrected populations (dashed bars, **Fig. S2B**) showed that, for the WT promoter, the non-pausing events arise mainly due to limited resolution; in contrast, for the ΔP promoter, the main reason is actually the presence of non-pausing RPs (**Fig. S2B**). The T+^6^G+^7^ (ntDNA) sequence therefore enforces pausing at ITC6 for ∼100% of ITCs, stabilizing the pre-translocated state arising from the clash between σ_3.2_ and the 5’-RNA end ^27,31^.

Finally, a fully double-stranded promoter (dsWT, **Fig. S1A**) did not modify the ITC6 pause exit rate both for ApA and ATP starting substrates (**Fig. S2C**), while the probability to reach the FS state during the first attempt on this promoter was also strongly decreased in the absence of a 5’-RNA end triphosphate (∼14% vs ∼58%, **Fig. S2D**), suggesting again that the 5’-RNA end triphosphate assists in the ITC6 pause exit.

### Weaker RNAP–promoter interactions promote cyclic scrunching/unscrunching

Our single-molecule reaction trajectories demonstrated (**Fig. 1**) that the transcription complexes paused at ITC6 may either resume RNA extension or cycle between stable paused states. The first apparent event on the cycling pathway is the isomerization of the PS promoter conformation to the US state. A major factor determining the partition of ITC6 to the productive or unproductive pathway could thus be the stability of the scrunched DNA conformation. To explore this hypothesis, we engineered several structural changes (**Fig. 3AB**), which alter important interactions between RNAP and nucleic acid components of the ITC (thus affecting scrunching), and characterized the effects on initial transcription.

**Fig. 3.**
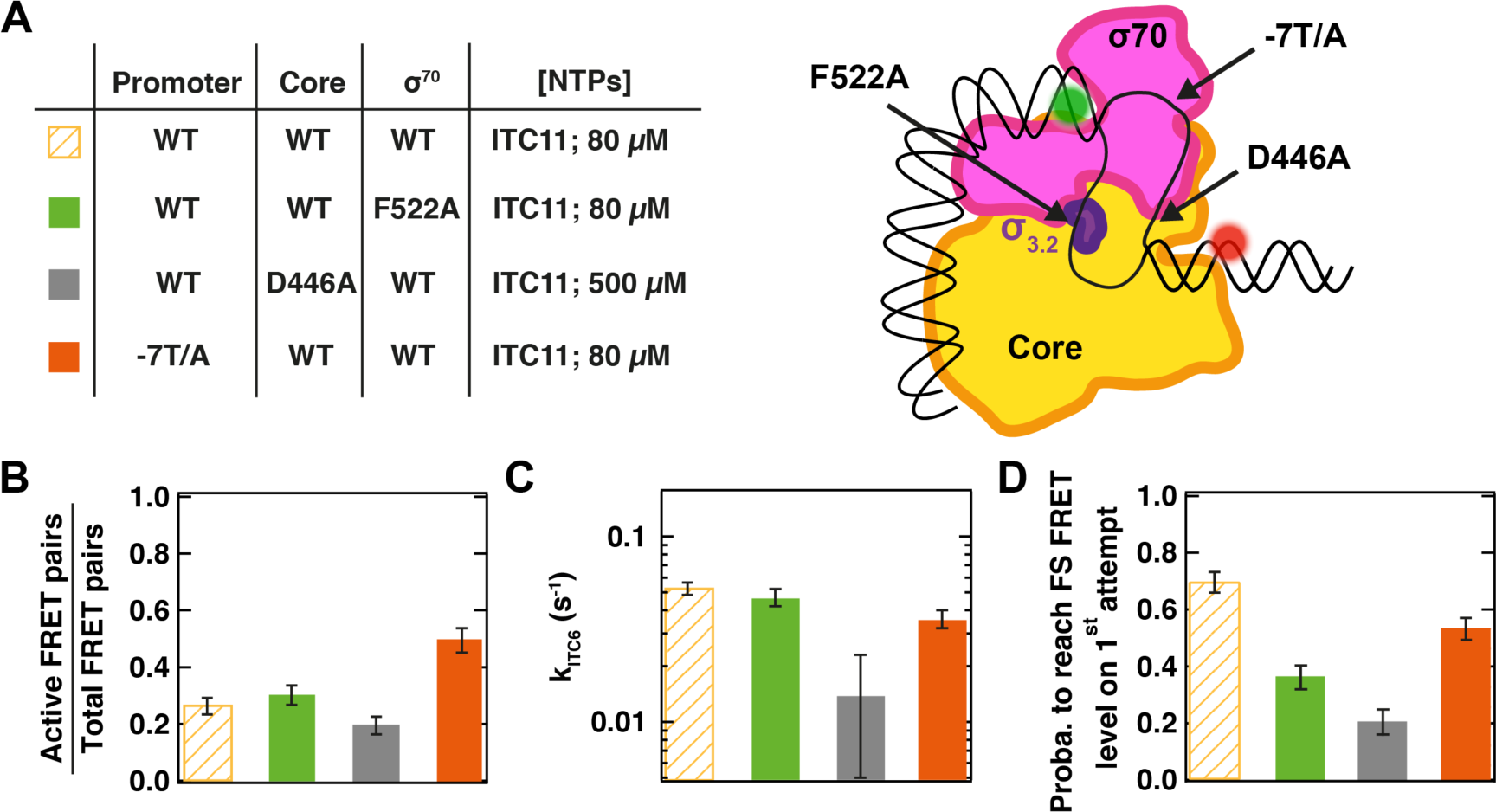
Core and σ^70^ mutants affect ITC6 pause exit probability. **(A)** Experimental conditions studied here and schematic of the different RP complex variants, all with ApA starting substrate. The same experiments have been performed with ATP starting substrate and are presented in **Fig. S2E-G**. **(B)** The fraction of transcriptionally active RPs, which displayed NTP-dependent E_FRET_ changes, of all surface-immobilised RPs. (see also, **Materials and Methods: FRET pair localization and detection**),in the experimental conditions described in (A). **(C)** ITC6 pause exit rate for the experimental conditions described in (A). **(D)** Probability to reach the FS FRET level on the 1^st^ attempt for the experimental conditions described in (A). Error bars are either one standard deviation from 1000 bootstraps procedure (C) or 95% confidence interval (B, D).

To establish the importance of interactions of σ_3.2_ with the template-strand DNA, we studied the F522A substitution in σ^70^, which eliminates an interaction between the −4 template DNA base and σ_3.2_ ^29^; this mutation has been shown to affect initial transcription, most notably by reducing the amount of transcripts shorter than 6-nt ^22^, and could therefore affect ITC6 pausing. We observed that the F522A σ^70^ derivative retained similar activity (**Fig. 3B, Fig. S2E**) and ITC6 pause exit rate (*k*_ITC6_) as the WT σ^70^ (**Fig. 3C, Fig. S2F**). Instead, the substitution significantly decreased the fraction of complexes exiting the pause on the first attempt from ∼70% to ∼37%, independently of the use of ApA (**Fig. 3D**) or ATP (**Fig. S2G**) as starting substrate. The weakening of σ_3.2_ interaction with the template-strand DNA thus destabilizes the PS promoter conformation and biases the paused ITC6 towards the scrunching/unscrunching pathway.

We next studied the effect of the β D446A RNAP substitution on ITC6 pausing. This mutation impairs the ‘G pocket’ in the RNAP core recognition element that specifically binds a guanine at ntDNA position +1 in the post-translocated state ^29^, strengthens the holoenzyme–promoter interaction ^29^ and helps to overcome a consensus elongation pause by stabilizing the RNAP at the post-translocated register ^33^. At the same time, these interactions stabilize a hairpin-depended pause ^36^. Notably, the post-translocated ITC6 on our WT promoter has a guanine in a position optimal for interacting with the G pocket (**Fig. 1A**). Our results demonstrate that the G pocket (in addition to being essential for forming an active ITC; **Fig. 3B, Fig. S2E**) facilitates pause exit, as from a consensus pause during elongation. Specifically, we observed ∼2-fold reduction in the pause exit rate (∼0.1 vs. ∼0.055 s^-1^, with ATP starting substrate **Fig. S2F**). We also observed up to 4-fold reduction in the fraction of complexes escaping the pause on the first attempt (∼20% vs ∼80% for ApA starting substrate, WT promoter and 500 μM NTPs, see **Fig. 3D** and **Fig. 2C**, respectively). The decreased pause-exit rate for the βD446A RNAP suggests that the paused ITC6 is biased towards the pre-translocated state, similar to the consensus paused elongation complex ^33^.

To probe the effects of weakened interactions between σ region 2 and the −10 promoter element, we also replaced the consensus −7 thymine in the non-template DNA by an adenine (-7T/A, **Fig. S1A**); σ specifically unstacks and inserts the thymine into a deep pocket during RP_o_ formation ^29,37,38^. Our experiments using −7T/A promoter show only small changes in the ITC6 pause exit rate and the fraction of complexes exiting the ITC6 pause on the first attempt (**Fig. 3BC**), showing that this interaction is not affecting significantly this phase of initial transcription.

#### Complexes undergoing cyclic unscrunching/scrunching are inactive for many minutes

We then quantitatively analyzed the ITCs that first pause at ITC6, and then perform cyclic unscrunching/scrunching. As seen in **Fig. 1D**, these complexes may cycle multiple times between the PS and US states until they reach the FS FRET level. Since cycling often lasted tens or even hundreds of seconds, many of the analyzed trajectories were interrupted by dye bleaching before the RP reached the FS state (**Fig. 4A**). For the cycling population, we generated probability density distributions for the dwell times in PS (Δt_PS_) and US (Δt_US_) states (**Fig. 4BC**). Both PS and US distributions showed a similar trend, with dwell times varying from ∼0.4 s to ∼200 s (**Fig. 4BC**). Using a MLE fitting routine (**Materials and Methods**), we found that the distributions were fitted well by a two-exponential probability distribution (solid lines, **Fig. 4BC;** dashed lines depict a single-exponential function). Our fit can thus define the exit rates *k*_1_ and *k*_2_ for both PS and US states, and the probability *P*(k_1_) to exit the US or PS state with the pause exit rate *k*_1_ (**Fig. S2E-G**; the probability to exit a state with the rate *k*_2_ is given by 1-*P*(k_1_)).

**Fig. 4.**
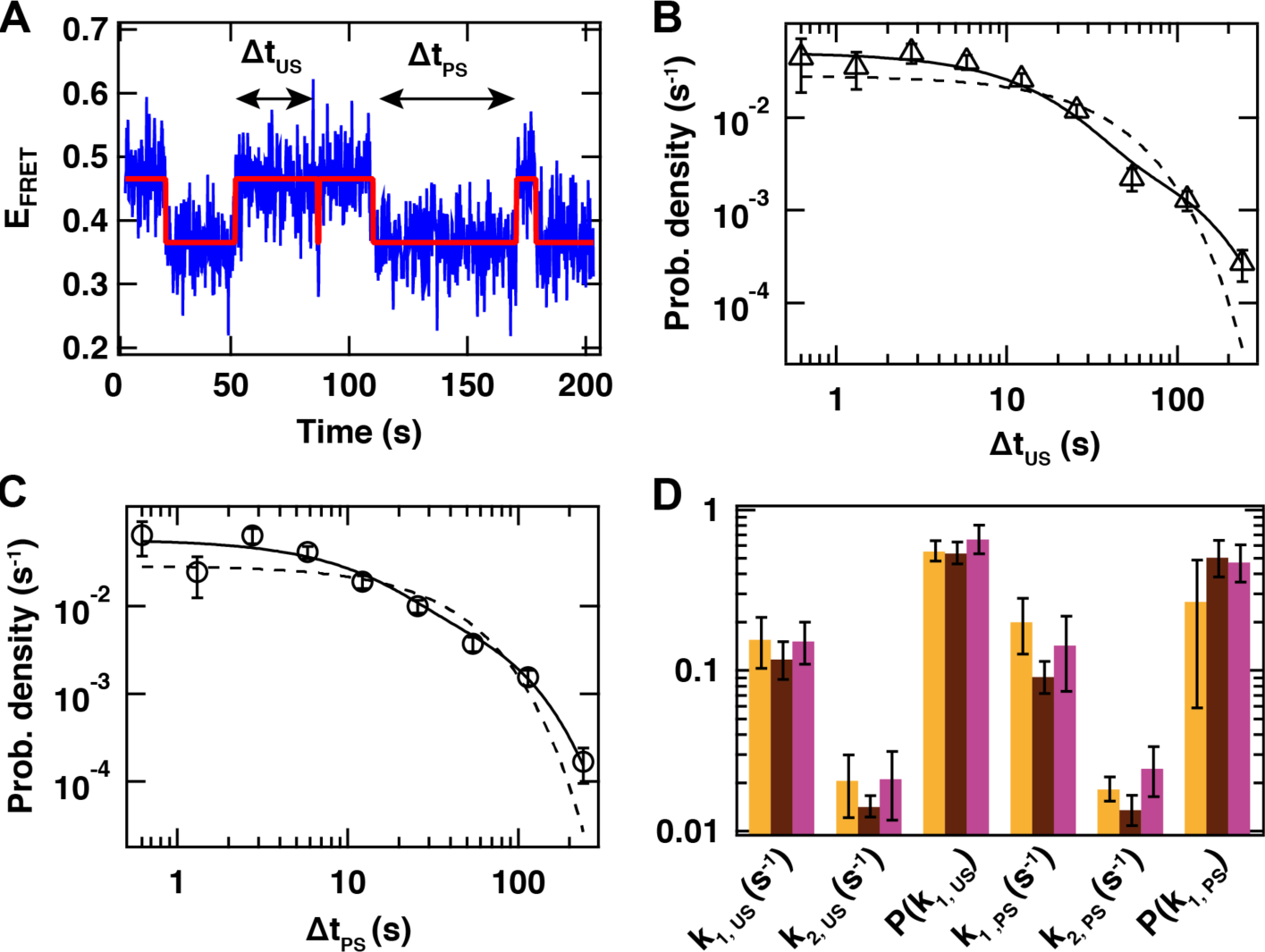
RP complexes that do not reach the FS FRET level alternate between US and PS FRET levels for long period of time. **(A)** Typical E_FRET_ trace where the RP alternates between unscrunched (US) and partly scrunched (PS) DNA promoter FRET levels. The red solid line represents the FRET levels extracted from empirical Bayesian probability Hidden Markov Model. We collect the dwell times Δt for each FRET level. Experimental condition: WT DNA promoter; NTP start: ApA; NTP: U/G/ATP 80 μM. **(B, C)** Probability density distribution of the dwell times Δt_US_ and Δt_PS_, respectively, for the experimental conditions described in (A), with its single and double exponential MLE fit (dashed and solid line, respectively, **Materials and Methods** for MLE fit procedure). **(D)** Average for all the NTP concentration (**Fig. S2E-G**) of the double exponential MLE parameters (k_1_ and k_2_ exit rates and P probability of being in the exponential with k_1_ exit rate) for US and PS FRET states averaged over all NTP concentrations, for the conditions described in **Fig. 2A** and **Table S1**. Color scheme as in **Fig. 2**.

We applied this analysis to our results from WT promoter reactions initiated with ApA or ATP, and the ΔP promoter initiated with ApA (**Fig. 2B**). We did not include the ATP-initiated ΔP promoter results, since most complexes exited the ITC6 pause directly to the FS state (**Fig. 2C**). We first noted that the exit rates *k*_1_ and *k*_2_, as well as the *P*(k_1_) probabilities of PS and US states, remained fairly constant in all used NTP concentrations (**Fig. S2H-J**). We observed a single exception with the ITC on the ApA-initiated WT promoter, which showed a decreased probability *P*(k_1, PS_) at higher NTP concentrations (right panel, **Fig. S2H**). To further improve our accuracy, we averaged the kinetic parameters over all used NTP concentrations. The US and PS states had practically identical kinetics, with the average values being *k*_1_∼0.15 s^-1^, *k*_2_∼0.02 s^-1^ and *P*(k_1_)∼0.6. Notably, these values were also independent of the NTP subset used (allowing maximal transcript lengths 7 or 11), the nature of the RNA 5’-end, the presence of pause motif in the transcribed sequence, the presence of the σ^70^ F522A and β D446A mutations, the fully double-stranded structure of the promoter, or the presence of the −7T/A substitution in the −10 element (**Fig. 4D, S2KLM)**.

The remarkable insensitivity of the kinetics of the unscrunching/scrunching pathway to the tested parameters, and in particular to the NTP concentration, suggests the complexes that enter unscrunching/scrunching pathway are catalytically inactive, until reentering the productive pathway to produce an ITC11 transcript (**Fig. 1D**). Indeed, if this pathway was catalytically competent, the exit rates *k*_1_ and *k*_2_ would have been sensitive to the NTP concentration, and would have therefore followed a Michaelis-Menten description. As evidenced from the slow unscrunching/scrunching rate constants, any complex embarking on the unscrunching/scrunching pathway very significantly delays the clearing of the promoter for the next cycle of transcription initiation, which potentially decreases the level of expression of the downstream gene.

### DNA unscrunching does not necessarily lead to abortive RNA release and re-initiation

The discovery of extensive cycles of unscrunching and scrunching during initial transcription raises intriguing questions about its relation with abortive initiation. Does each unscrunching event lead to the release of nascent RNA (**Fig. 5A**, left panel)? Is the subsequent re-isomerization to scrunched state driven by the synthesis of new RNA? Could the RNA be maintained in the complex upon unscrunching (**Fig. 5A**, right panel)? To address these questions, we performed experiments (**Fig. 5B**) in which we allowed RNA synthesis up to ITC11 (**Fig. 5EG**) or ITC7 (**Fig. S3A**) for ∼10 s, washed the surface extensively to remove NTPs, and re-imaged the surface-bound complexes. To our surprise, we observed many complexes displaying unscrunching/scrunching activity in the *absence* of NTPs; The percentage of complexes cyclically unscrunching/scrunching in the absence of NTPs was ∼28% and ∼18% for the WT promoter initiated with ApA or ATP, respectively (**Fig. 5D**). These numbers should be compared to ∼27% (ApA) and ∼42% (ATP) of active FRET pairs in the presence of NTPs, respectively (**Fig. 5D**). This means that a large fraction of the cycling molecules in the presence of NTPs (potentially up to the entire population, in the case as ApA-initiated reactions) can be accounted for by cycling complexes that do not synthesize RNA. The scrunching/unscrunching cycling lasted for hundreds of seconds, being only limited by dye bleaching (**Fig. 5EG**). Consistent with the maximal RNA length, complexes pulsed with ITC7 NTP sampled only US and PS states (**Fig. S3A**) whereas complexes pulsed with ITC11 NTP could additionally occupy the FS state (**Fig. 5G**). Our results clearly establish that extended cycling in different scrunching states does thus not require active RNA synthesis.

**Fig. 5.**
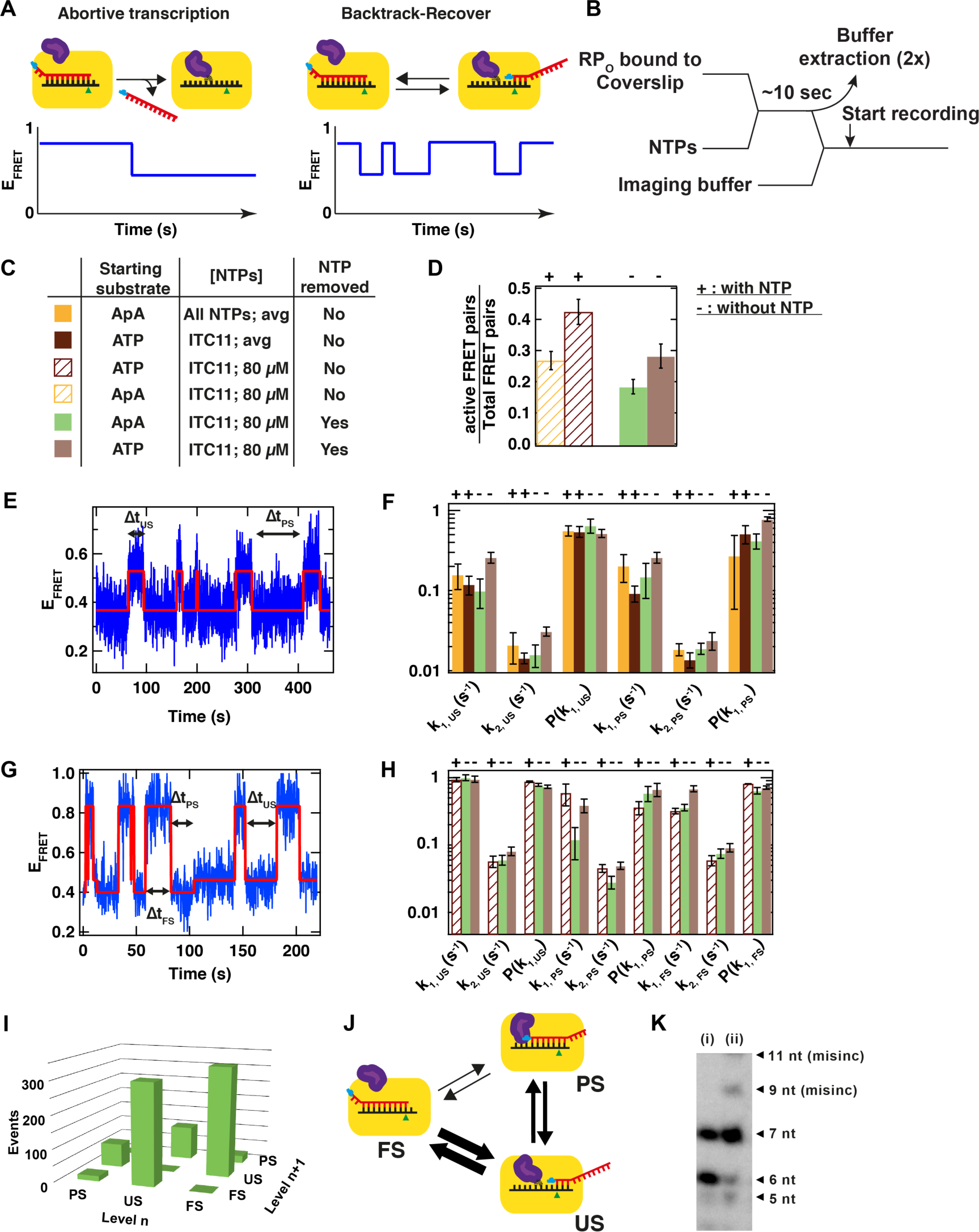
Post-RNA synthesis DNA promoter scrunching kinetics are independent of the presence of NTP. **(A)** Schematic of the RPs possible behavior and the corresponding FRET signal: on the left hand side, the RP releases the nascent transcript upon DNA unscrunching, displaying a single step kinetics in the smFRET trace; on the right hand side, the template DNA/RNA hybrid alternates between a pre-translocated register and a backtrack position upon the FS→US transition, while retaining the RNA. **(B)** Schematic of the experiment. **(C)** Table presenting the experimental variables probed in **Fig. 5**. Note that the data represented in brown and yellow are from **Fig. 2** and are averaged (avg) values for all the concentration of NTP probed in these experiments. Here, we used the WT DNA promoter (**Fig. S1**) and wild-type holoenzyme (**Materials and Methods**). **(D)** Ratio of the numbers of active FRET pairs, i.e. that display scrunching/unscrunching cycles, over the total number of FRET pairs after selection (**Materials and Methods: FRET pair localization and detection**). The + and - above the bars indicate the presence and the absence, respectively, of NTPs during the experiment. An identical notation is used in (F) and (H). **(E)** Experimental FRET trace after NTP removal (B) showing an RP complex alternating between US and PS, but not in the FS FRET level. The experimental conditions used for the acquisition of this trace correspond to the light green color code in (C). **(F)** Scrunching kinetics (k_1_, k_2_ and P(k_1_)) extracted from a double-exponential MLE fit analysis of the Δt_US_ and Δt_PS_ distributions from the traces alike (E). **(G)** Experimental FRET trace after NTP removal (B) showing an RP complex alternating between US, PS and FS FRET level. **(H)** Scrunching kinetics (k_1_, k_2_ and P(k_1_)) extracted from a double-exponential MLE fit analysis of the Δt_US_, Δt_PS_ and Δt_FS_ distributions from the traces alike (G) for the conditions described in (C). **(I)** 3D histogram showing the number of transition between two different FRET levels for two consecutive dwell times n and n+1 for the traces acquired as described in (C). **(J)** Schematic resuming the results of (H); the thicker the arrow, the more likely the transition. **(K)** Transcript retention experiment using magnetic bead attached RP complexes incubated in ITC7 conditions and rinsed (i), then restarted after 15 sec. by chasing 80 μM GTP (ii) (**Supplementary Protocol**). The 9mer and 11mer originated from GTP misincorporations.

We quantitatively analyzed the kinetics of cyclic unscrunching/scrunching for complexes pulsed with ITC11 NTP, and identified two subpopulations: the first cycled between US and PS FRET levels only (**Fig. 5E**), and the second cycled between US, PS and FS FRET levels (**Fig. 5G**). The US/PS subpopulation included ∼50% (ApA starting substrate) or ∼40% (ATP starting substrate) of all cycling molecules, respectively (**Table S1**). The US/PS and US/PS/FS subpopulations most likely represent the ITCs which, at the moment of NTP withdrawal, had synthesized 6- and 11-nt RNAs, respectively. The cyclic unscrunching/scrunching kinetics of US/PS subgroup did not differ significantly from that observed in the continuous presence of NTP, i.e. *k*_1_∼0.15 s^-1^, *k*_2_∼0.02 s^-1^ and *P*(k_1_)∼0.6 (**Fig. 5F**). Interestingly, the US/PS/FS subgroup sampled all scrunching states almost an order of magnitude faster compared to the US/PS (i.e. *k*_1_∼1 s^-1^, *k*_2_∼0.06 s^-1^ and *P*(k_1_)∼0.7, **Fig. 5H**). Using ATP or ApA for initiation did not significantly affect the kinetic parameters of the US/PS/FS FRET states (**Fig. 5FH**).

Close inspection of the trajectories belonging to the US/PS/FS subgroup revealed that the two most frequently encountered state transitions were FS→US and its reversal US→FS (**Fig. 5I** and **Fig. S3F**); this was also the case in the continuous presence of NTP (**Fig. S3G**). The US→PS and PS→US transitions were about 4-fold less frequent, whereas PS→FS or FS→PS transitions were only rarely observed. This data clearly indicate that RPs engaged in the unscrunching/scrunching pathway do not share the same linear US→PS→FS reaction coordinate of ITCs engaged in productive transcription (**Fig. 5J**). We also note the absence of any temporal correlation between two successive state dwell times (dt_n_ and dt_n+1_), independent of the scrunching state they originate from (right hand side, **Fig. S3BCD**), which shows that the transition from one state to the next is memory-less, i.e., the scrunching magnitude of the preceding state has no effect on the timescale of the transition of the following state.

### Paused ITC may undergo abortive initiation or stably trap RNA

Our FRET assay monitors the conformation of the promoter DNA and thus does not provide a direct readout for the presence of RNA in the ITCs. Since pulsed RNA synthesis was required to generate ITCs that cycle for several minutes between scrunched states, we assumed that these ITCs retain the nascent RNA in the transcription bubble. The assumption generates two testable hypotheses: first, RNA is slowly released from NTP-deprived ITCs; second, any RNAs retained in ITCs are extendable upon NTP reintroduction.

To determine the profile and time-dependence of RNA release from ITCs, we immobilized biotinylated RP_O_ complexes to streptavidin-coated magnetic beads. The complexes were pulsed for 10 s with the ITC7 NTP subset (containing σ−^32^P-UTP), pulled down, washed and immersed into NTP-free reaction buffer; beads and supernatant were then analyzed at specified times to obtain the time-dependent profile of retained and released RNAs (**Fig. S4AB**). Our results showed that the RNA-release kinetics was strikingly biphasic: many ITCs released their RNA within the first 2 min, the release being almost quantitative for the shortest RNAs (∼95% of 3–4-mers) and less efficient for 5-, 6-, and 7-nt RNAs (45, 80, and 80%, respectively; **Fig. S4B**). After the rapid initial phase, the amount of released 6- or 7-nt RNA increased only marginally. After 15 min, still ∼20% of 6–7-nt RNA remained bound in the ITCs. This amount is 2-fold lower than what we measured in similar NTP-pulsed single-molecule experiments, where most of the active ITCs were sampling the unscrunching/scrunching states for several minutes (**Fig. 5D**).

To probe whether the stalled ITCs retaining 6-nt RNA for an extended period of time can resume active transcription, we chased the immobilized and washed ITCs with the next incoming nucleotide (GTP). We observed that the 6-nt RNA became converted quantitatively to 7-nt RNA (**Fig. 5K;** longer products appear due to mis-incorporation), indicating that the ITCs both retain the nascent RNA in the transcription bubble, and can access the catalytically active conformation.

In summary, the biochemical analysis revealed two populations of stalled ITCs: 70–80% of ITCs that enter the abortive initiation pathway (rapidly releasing the nascent RNAs) and 20– 30% of ITCs that retain 6–7-nt RNA products and catalytic competence for tens of minutes after NTP depletion. A possible cause of the two distinct ITC behaviors could relate to the open complex or σ_3.2_ conformations, which undergo extensive remodeling during initial transcription (see *Discussion*). These results clearly show that abortive RNA-release and transcription reinitiation are not obligatory upon DNA promoter unscrunching. The nascent RNA can be stably trapped within the cyclically unscrunching/scrunching RP complex, until being eventually elongated.

## Discussion

In this study, we employed a refined single-molecule FRET assay to quantitatively dissect the reaction pathway and kinetics of the initially transcribing complexes on the *lac* promoter. Our unique FRET sensor helped us to observe directly and with high contrast the entrance and exit from initiation pausing and allowed us to disentangle the complex network of catalytic and non-catalytic events during initial transcription, and to examine the role of the σ_3.2_ region, the nature of pausing, and pausing-related conformational changes such as scrunching/unscrunching in the presence and absence of RNA release.

### σ_3.2_ represents a translocation barrier during initial transcription

Two aspects of our current results and our previous work ^27^ confirmed the presence of a rate-limiting pause during initial transcription on *lac* promoter: first, the biochemical analysis demonstrated accumulation of a 6-nt RNA product within the transcription complex; second, the single-molecule reaction trajectories demonstrated that nearly all RPs (80–90%) progressing to fully scrunched promoter state (ITC11 or All NTP) showed a ∼5–20 sec preceding dwell at the partially scrunched state, which we infer as ITC6. A protein element that was a good candidate for causing pausing was the σ_3.2_ region, that was shown by structural, biochemical and single-molecule biophysical studies to occlude the RNA-exit channel of RNAP and to form a barrier for the elongation of the nascent transcript past 5–6-nt ^12,19,20,27^ (**Fig. 1B**, **Fig. 6**). Consistent with these results, we recently showed that partial deletion of σ_3.2_ significantly diminished pausing at ITC6 ^27^.

**Fig. 6.**
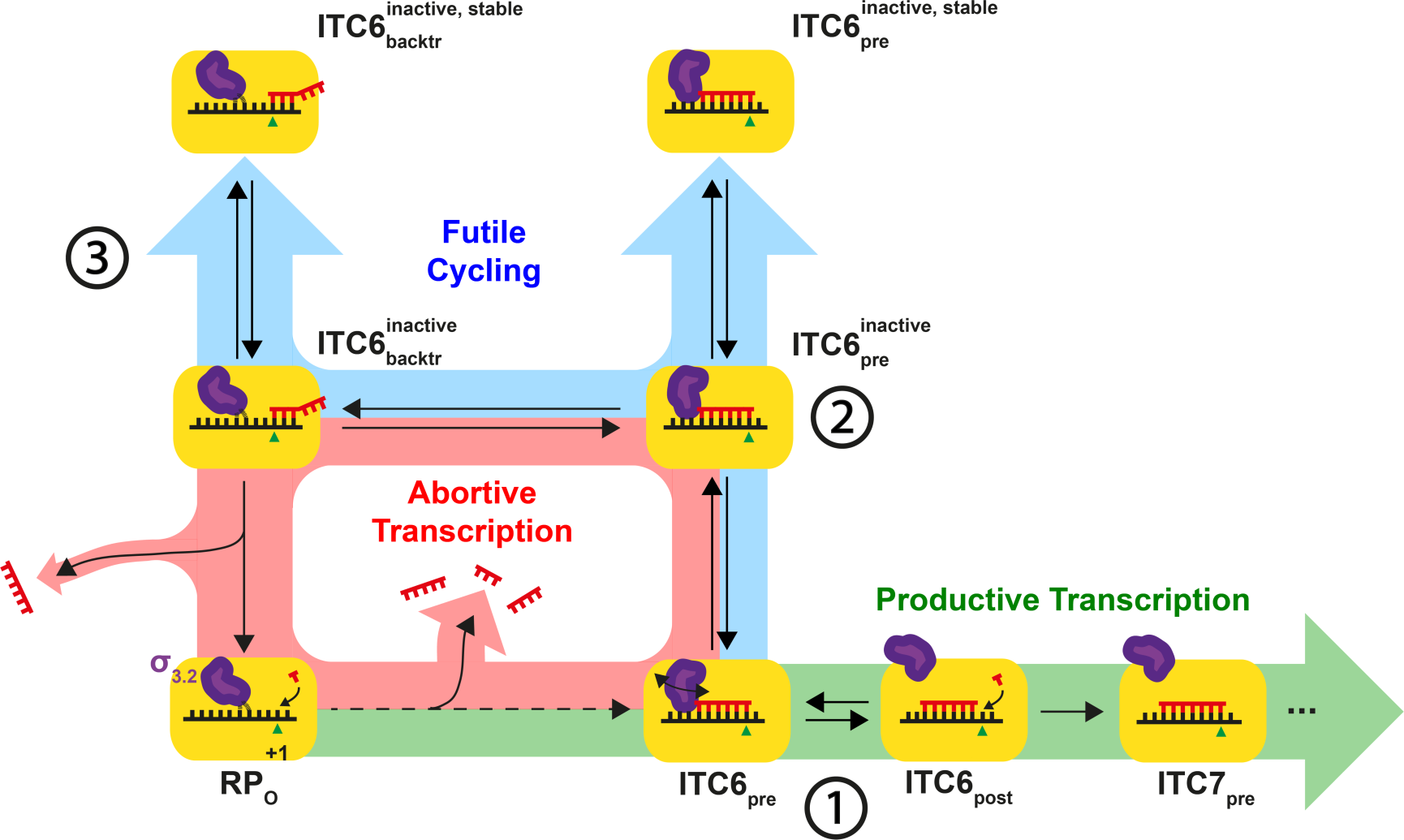
Model for initial bacterial transcription. The progress of initial transcription is illustrated by depicting RNAP (yellow block) at key points of the inferred mechanism. The mechanism includes three competing reaction pathways, which the ITC can embark on. Productive Transcription pathway (highlighted in green) results in promoter escape and synthesis of full-length RNA. Abortive Initiation pathway (highlighted in red) leads to the synthesis and dissociation of short RNA products. Futile Cycling (highlighted in blue) temporarily traps the ITC6 into catalytically inactive interconverting pre-translocated and backtracked states, respectively. Purple finger shows the different conformations of σ_3.2_. Green triangle marks the template base for the next incoming nucleotide in the active site of RNAP. Red and black strands represent the nascent RNA and template DNA, respectively. The numeration (1, 2 and 3) indicates the three significant molecular mechanisms described by the model: the initial barrier imposed by σ_3.2_ to the transcript elongation, the subsequent loss of catalytic conformation and the RNA-dependent reversible backtracking, respectively.

However, since the same σ_3.2_ derivative was associated with accelerated conformational dynamics in the open complex ^39^, a possibility existed that some of the inferred effects on pause kinetics were indirect (e.g. due to instability of the template strand conformation in the DNA binding cleft leading to increased abortive RNA-release and shortening of the ITC6 pause). Our new finding that the triphosphate moiety at the 5’-RNA end, which specifically interacts with the σ_3.2_ ^12,20^, both shortens the half-life of ITC6 pause and increases the probability of productive pause exit confirms the role of σ_3.2_ as a major pause determinant in initial transcription.

The 80–90% probability to enter the pause may reflect the presence of transcriptionally non-permissive (pausing RPs) and permissive (non-pausing RPs) σ_3.2_ conformations present in different ITC6 complexes. Based on the structural considerations ^12,20^ and the stage of initial transcription (Ref. ^27^; this study), the clash between σ_3.2_ and RNA 5’-end may hamper the movement of the template DNA and/or RNA to the post-translocated register after the addition of the 6^th^ nucleotide. A key mechanistic aspect of the ITC6 pause could thus be the stabilization of the pre-translocated state ^27,31^. We provide additional strong evidence in favor of this hypothesis by showing that βD446A RNAP (which de-stabilizes the post-translocated state due to the loss of the +1 ntDNA guanine interactions with a specific β pocket ^33^), displayed two-fold decreased ITC6 pause exit rate. However, because the pause exit rate did not strongly depend on the NTP concentration (increasing only from 0.2 to 0.3 s^-1^ when NTP was raised from 5 to 500 μM), the pause is *not* directly controlled by the thermodynamic equilibrium between the pre- and post-translocated states of ITC6. By similar reasoning, the pause is also not controlled by the catalytic rate of post-translocated ITC6. We thus postulate that the pause-controlling step is kinetic and involves relatively slow repositioning of the σ_3.2_ tip in a way that the barrier to forward translocation is removed. The highest observed pause exit rate on the promoter variant lacking the consensus pause motif (0.3 s^-1^ for the ΔP promoter, **Fig. 2B**) may reflect the rate of ITC6 pre→post translocation that we suggest to be controlled by σ_3.2_ repositioning. This rate would signify dramatically longer dwell (∼265-fold: 2.3 s vs. 8.7 ms) in the pre-translocated state in ITC6 compared to the dwell in the pre-translocated state during unimpeded cycle of transcription elongation ^40^. Importantly, several studies of pausing during transcription elongation have shown the predisposition of the pre-translocated RNAP to isomerize into a catalytically inactive off-pathway state, known as the elemental pause ^2,33,34,41^. The σ_3.2_-dependent translocation barrier encountered during initial transcription may thus act, by accumulating the pre-translocated ITC6, to increase the probability to isomerize into an elemental pause-like state.

### Initial transcription pause involves elemental pause–like states

The finding that the Δσ_3.2_ mutation reduced but did not completely abolish pausing at ITC6 indicated that σ_3.2_ is not the only determinant of initial transcription pausing ^27^. Recent work elucidated a consensus sequence that dramatically increased the probability of transcription elongation complex to enter the consensus pause ^33,34^. The most strongly conserved part of the consensus pause motif is a pyrimidine-guanine (Y)/G at position −1/+1 relative to the 3’- end of the transcript, which may cause the template strand to isomerize in the pause complex such that the template base becomes inaccessible to the incoming NTP ^2,41^. A consensus pause motif is indeed encountered at ITC6 ^31^ and, importantly, substitution of the motif increased both the pause exit rate and the probability to exit the pause towards ITC7; these results are consistent with biochemical studies of many promoters, including the *lac* promoter ^31^. The exit rate from the ITC6 pause (∼0.3 s^-1^) is similar to the exit rate from consensus elongation pause (∼0.5 s^-1^) ^34^. Overall, it appears that the first events leading to a pause during initiation and elongation phases of transcription are similar: an energetic (transcribed sequence in elongation) or physical (σ_3.2_ in initial transcription) barrier to translocation delays RNAP in the pre-translocated register ^31^ from where the protein can, with sequence-dependent efficiency, branch-off to a catalytically inactive elemental pause state (**Fig. 6**).

### Backtracking leads to long-lived paused states

While the entry of ITC6 into the elemental pause was nearly obligatory (80–90% of trajectories showed the pause, **Fig. S2B**), a significant fraction (∼20% at saturating NTP concentration, **Fig. 2D**) of the RNAP complexes did not exit this pause on the first attempt, but instead embarked on another reaction pathway involving cyclic unscrunching/scrunching events. Provided that the initially transcribing RNAP remains tightly anchored at the promoter, an unscrunching event results in partial relaxation and reannealing of the downstream DNA that was pulled into RNAP in the scrunched state ^10^. Another important consequence of unscrunching is the displacement of the 3’-RNA end from the active site to the NTP-entry channel and, hence, catalytic inactivation of the complex. The unscrunching therefore mechanistically resembles the backtracking of transcription elongation complex and effectively, similar to some elongation pauses ^42^, serves to isomerize the ITC6 at the elemental pause to long-lived inactive states. The probability to enter the unscrunching/scrunching pathway inversely correlated with NTP concentration suggesting that, when the cellular NTP pool is low, the unscrunching/scrunching mechanism can efficiently inhibit promoter escape and, hence, decrease transcript levels. Furthermore, weakening the holoenzyme interactions with the template DNA (by F552A substitution in σ_3.2_) or non-template DNA (D446A substitution in β) favored the partitioning of ITC6 into the unscrunching/scrunching pathway (**Fig 3D**, **Fig. S2G**). This finding may imply that the promoter and initially transcribed sequences, interacting with the holoenzyme most tightly, encode efficient promoter-escape kinetics because they disfavor ITC partitioning into the non-productive unscrunching/scrunching pathway. Consistently, Record and co-workers recently reported the correlation of stronger holoenzyme–discriminator (promoter sequence between the −10 element and transcription start site) interaction with the production of longer abortive RNAs, while having a higher promoter escape efficiency ^43^.

Similar backtracked/unscrunched initially transcribing complexes were recently identified in a magnetic tweezers assay ^28^. Both that study and our current work described the US state kinetics with a double-exponential distribution, with the longest-lived one being the backtracked complexes. Lerner et al. ^28^ reported backtracked complex lifetimes two orders of magnitude longer than we find in our study. However, the difference can be easily explained as the magnetic tweezers assay of Lerner et al. is based on the transcription-dependent changes in the supercoil density of a positively supercoiled DNA. Essentially, RNAP activity, e.g. open complex formation and RNA synthesis, generates plectonemes (“DNA loops”) ^11,28,44,45^. However, the energetic cost of plectoneme formation ^46-49^ biases the transcribing complex towards less extensive scrunching, increasing the lifetime of the backtracked/unscrunched state. Our FRET assay, in contrast, employed non-supercoiled DNA and reported a shorter lifetime of the backtracked/unscrunched state. In addition, the symmetry in the dynamics of the US and the PS states, together with its independence on catalytic activity or the nature of the interactions between the RNAP and the promoter, suggest that the transition between the states described by two exponentials, i.e. described by exit rates k_1_ and k_2_, (**Fig. 4 and Fig. S2H-M**) originates from a conformational change within the holoenzyme that precludes catalysis but does not preclude the dynamics of the hybrid, i.e. scrunching/unscrunching cycle.

### RNA release and subsequent re-initiation is not obligatory upon DNA unscrunching

Previous single-molecule studies assumed a direct link between unscrunching and abortive transcription ^11,17,27,28^. However, those studies focused on the DNA conformation and did not evaluate the presence of RNA in the transcription complexes upon unscrunching Our data demonstrated that brief pulsing of open complexes with NTP resulted in a population of ITCs that kept on cycling between US/PS/FS states for an extended period of time. We also showed that, if not immediately released, the transcripts remained stably attached to the RNAP for at least 45 min. Backtracking to the US state is expected to shorten the template DNA–RNA hybrid to ≤5 bp, and should reduce the hybrid lifetime. However, the possibility of a short hybrid that locks the transcript into the DNA binding cleft is supported by the observation of a 4-nt RNA bound to a bacterial open promoter complex in crystals ^29,37^. Furthermore, the backtracked RNA potentially forms interactions in the NTP-entry channel, as observed in the yeast RNA polymerase II ^50^. Taken together, the stability of the short hybrid within the complex and the positive interactions between the RNA and the NTP entrance channel support the absence of transcript release upon promoter unscrunching. We note that the presence of the GreA and GreB factors *in vivo* likely shortens the lifetime of the backtracked US state by cleaving the protruding 3’-RNA end of backtracked complexes ^51^, as observed by Lerner et al. in *in vitro* single-molecule experiments ^28^. Unfortunately, we could not evaluate the effects of Gre-factors with our single-molecule FRET assay because their binding to RP caused extensive flickering of the Cy3B dye.

Recent work ^43^ have also noted that RP_O_ complexes on λP_R_ and T7A1 promoters were divided into two populations upon NTP addition: a first population (30-45% of all complexes) that rapidly (within 10 sec) synthesized long RNA, i.e. longer than 10 nucleotides, quickly, i.e. within 10 sec, and represented 30-45% of the total RP population; and a second population that was stalled in early ITC, i.e. shorter than ITC10, and that released RNA slowly, similarly to moribund complexes ^52^. We propose that these two populations, i.e. the population producing quickly long RNAs and the moribund complexes, are consistent with the two populations we described here, i.e. the RP complexes that exited the ITC6 pause on the first attempt (**Fig. 1C**), and the population that entered the cyclic unscrunching/scrunching state from the ITC6 pause (**Fig. 1D**), respectively. Henderson et al. reported that the moribund complexes on their promoters and under their conditions were not able to elongate their transcripts ^43^. In contrast, our complexes are able to extend the retained RNA (**Fig. 5K**), and could enter the cyclic unscrunching/scrunching state multiple times before elongating the transcript, as a function of the NTP concentration (**Fig. 1D, Fig. 2C**). This conclusion is further supported by the biphasic activity also observed with purified moribund complexes ^52,53^, demonstrating that moribund complexes have entered a catalytically incompetent state, which is eventually exited towards productive synthesis. Supporting our conclusion on the backtracked nature of the US state, Shimamoto and co-workers have also observed that the moribund complexes were reactivated by GreA ^53^. They also showed that part of the moribund complexes are converted into dead-end complexes on λP_R_ promoter, i.e. that could not show any catalytic activity anymore, while trapping a short transcript ^52^. Interestingly, their gels showed that a large amount of 9-mer transcript was produced, despite the presence of all NTPs, which correlated with the presence of a (YG) sequence on the non-template promoter DNA at the +9 position. We showed here that such a (YG) sequence increased the probability to enter the cyclic unscrunching/scrunching pathway (**Fig. 2C**). We suggest that the dead-end complexes observed on the λP_R_ promoter are the consequence of multiple successive entries into the cyclic unscrunching/scrunching pathway, and therefore appeared inactivated. Similarly, it is likely that the complexes observed by Henderson et al. ^43^ have not yet recovered from the cyclic unscrunching/scrunching state. Nevertheless, our data clearly demonstrate that the unscrunching of the promoter DNA is not linked with obligatory abortive transcription and the release of RNA. Instead, off-pathway ITCs can sample different scrunched conformations and eventually resume productive RNA synthesis. Such conformational changes have also been proposed for promoter-proximal paused complexes ^54^.

### A model for initial transcription

We summarize our findings in a new kinetic model of the transition to productive transcription (**Fig. 6**). In the RP_O_ complex, σ_3.2_ interacts with and stabilises the melted template DNA strand in the DNA binding cleft of RNAP. Upon NTP addition, RNAP engages RNA synthesis producing rapidly either abortive 2–5-mer RNAs or ITC6. In ITC6 stage, σ_3.2_ becomes a barrier to the 5’-end of RNA stabilising the pre-translocated ITC6 – the key intermediate of our model that can branch to two competing reaction pathways. The fraction of ITC6 that remain on the productive pathway eventually overcome the barrier for translocation, possibly via rate-limiting retraction of the σ_3.2_ tip towards the RNA exit channel, resuming RNA synthesis. The pre-translocated ITC6 may also embark on an alternative off-pathway by isomerisation into the initial paused conformation (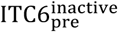 in **Fig. 6**). The paused state is further stabilised by the cyclic unscrunching/scrunching (or backtracking/forward-translocation) events depicted as “Futile cycling” in **Fig. 6**, which does not involve RNA synthesis or RNA release. The fact that the dwell-time distributions of both the PS and US states were double-exponential suggest that they can be further stabilized into long-lived PS 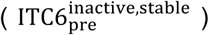 and US 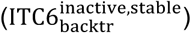 states, respectively. These states can retain the transcript for >45 min (**Fig. S4**) and ultimately return to the productive pathway and escape the ITC6 pause. The model we present here contains three significant molecular mechanisms, i.e. the initial barrier imposed by σ_3.2_ to the transcript elongation (1 in **Fig. 6**), the subsequent loss of catalytic conformation (2 in **Fig. 6**) and the RNA-dependent reversible backtracking (3 in **Fig. 6**), which potentiate, initiate and amplify the pause encountered by the initially transcribing bacterial RNAP, respectively.

### Biological significance

The dependence of entry and recovery from the pause states as a function of initially transcribed sequence ^31^ implies wide variation in the kinetics of initial transcription across the bacterial promoter sequence space. On the *lac* and similar promoters, the molecular mechanism of pausing sensitizes the efficiency of promoter escape to NTP concentration, potentially trapping the RNAP to the promoter in a “ready-to-fire” or “poised” mode until improved growth conditions lead to the replenishing of cellular NTP pool ^55-57^. The trapping of poised RNAPs at or near the promoter thus emerges as a common transcription regulation strategy achievable by different molecular mechanisms. For example, the σ^54^–RNAP holoenzyme forms an inactive, stable closed complex in bacteria ^58^, whereas negative elongation factors cause RNAP to stall within 20–60 bases downstream of the transcription start site in many metazoan genes ^59^. In all cases, inhibited RNAPs are ready-to-fire when activating signals arrive from relevant signal-transduction cascades thus bypassing potentially slow or stochastic steps in promoter search, binding, melting, and activation.

## Acknowledgements

We thank Dr. Francesco Pedaci and Tao Ju Cui for carefully reading the manuscript. We also thank Dr. Jan Lipfert for discussion.

## Author contributions

DD and ANK designed the research; DD designed, performed and analyzed the single-molecule experiments the single-molecule experiments; MK made the DNA promoter constructs; DD and JJWB designed and implemented the custom analysis routines; DLVB performed and analyzed the gel electrophoresis experiments; ZM and KB provided the σ70 mutant; IP and AK provided the core RNAP mutant; DD, AMM and ANK discussed the data; DD, AMM and MD discussed the model; DD, AMM, and ANK wrote the article.

## Materials and Methods

### Description of glass coverslips preparation for single-molecule experiments

Borosilicate glass coverslips (#1.5 MenzelGlÄzer, Germany) were sonicated for thirty minutes in a 2% (V/V) solution of Hellmanex III (Helma Analytics, Germany)/deionized water. After being thoroughly rinsed with deionized water, the coverslips were dried, disposed into a plasma cleaner (Harrick Plasma, NY, USA) and exposed to a nitrogen plasma for thirty minutes. The coverslips are subsequently immerged into a 1% (V/V) solution of Vectabond (product code #SP-1800, Vector Labs, CA, USA)/acetone for 10 minutes. The coverslips were then rinsed in deionized water and dried with a stream of nitrogen gas. After disposing a silicone gasket (#103280, Grace Bio-Labs, OR, USA) on each coverslip, each well was filled with 20 μl of a 100 mg/ml solution of methoxy-PEG (5kDa)-SVA/ biotin-PEG (5kDa)-SC (2.5% (w/w) (Laysan Bio, AL, USA) in 50 mM MOPS-KOH buffer, pH 7.5 The wells were incubated for ∼1.5 hours and thoroughly rinsed with a 1x phosphate buffered saline (Sigma Aldrich, UK) solution. The coverslips were stored at 4°C up to two weeks before use.

### Protein immobilization protocol

The pegylated coverslips were incubated for ∼10 minutes with a solution 0.5 mg/ml of Neutravidin (#31050,Sigma Aldrich, UK) in 0.5x PBS and subsequently rinsed with 1x PBS. Preceding observation on the microscope, the coverslips were incubated for ten minutes with a 3% (V/V) solution of Penta•His biotin conjugate antibody (#34440, QIAGEN, UK) in reaction buffer (40 mM HEPES buffer pH 7.3 (ThermoFisher Scientific, UK), 100 mM potassium glutamate, 10 mM MgCl_2_, 1 mM dithiothreitol (DTT), 1 mM cysteamine hydrochloride, 5% glycerol (V/V), 0.5 g/l BSA) and subsequently rinsed with reaction buffer. After adjusting the coverslips on the microscope, 100 pM of His-tagged protein-DNA complex, e.g. RPo, was incubated in the observed well until the desired density of molecules on the coverslips surface was reached, followed by one-step rinsing with reaction buffer.

### Core RNA polymerase and σ^70^ preparations

The expression and purification of the core bacterial RNA polymerase have previously been described in Ref. ^60^. The expression and purification of the wild-type σ^70^ have been previously described in Ref. ^21^.

### Holo-RNAP and RP_O_ assembly description

0.5 μM of VS10 core RNAP was mixed with 0.6 μM of σ^70^ in 20mM Tris-HCl pH 7.9, 150 mM NaCl, 0.1 mM EDTA, 50% (V/V) Glycerol and 0.1 mM DTT and incubated at 30°C for 30 minutes. The resulting holo-complex was stored at −20°C.

5 nM of holo-complex was mixed with 2.5 nM of DNA promoter in reaction buffer and incubated for 10 minutes at 37°C to form the RPo complex.

### DNA constructs preparation

The DNA constructs preparation has previously been described in detail in the **Supplementary Protocols**.

### Microscope and single-molecule experiments description

The single-molecule TIRF microscope for FRET experiments has been previously described in Ref. ^61^. Shortly, the 532 nm, donor laser excitation, and the 642 nm wavelength laser beams (donor laser excitation and acceptor laser excitation, respectively) were focused in the back focal plane of an oil immersion objective (Olympus, N.A. 1.4) and illuminates alternatively the field of view, i.e. ALEX mode ^62,63^. The TIRF-reflected beams were directed toward a position sensor to control the objective focal plane distance to the sample at a fixed position (MS-2000 stage, ASI, OR, USA). The photons resulting from the de-excitation of the dyes molecules, i.e. fluorescence, were separated from the excitation laser beams with a dichroic mirror and spectrally splitted in two channels, e.g. donor and acceptor that are imaged on the same EMCCD camera (iXon, Andor, Irlande). For 100 ms ALEX illumination, i.e. 200 ms frame time acquisition, the laser power measured preceding the dichroic mirror is ∼0.4 mW for donor excitation laser and ∼0.09 mW for the acceptor excitation laser. For 40 ms ALEX illumination (only used to acquire the data with the ΔP promoter and ATP starting substrate experimental condition), i.e. 80 ms frame time, the laser power measured preceding the dichroic mirror is ∼1 mW for donor excitation laser and ∼0.25 mW for the acceptor excitation laser.

The imaging buffer contained the reaction buffer completed with 1 mM Trolox, 1 mM COT, 1% (w/V) glucose, 0.4 μg/ml of catalase and 1 mg/ml of glucose oxidase (Sigma Aldrich). The catalase and the glucose oxidase were pre-mixed together in a solution of 50 mM KCl and 50 mM Tris-OAc buffer pH 7.3 at 100x concentration ^64,65^.

The data were acquired after immobilization of the RPo complex to the surface. After ∼200 frames (∼20 s) the imaging buffer is spiked with a 12.5x NTP solution and the reaction is observed for the remaining ∼5800 frames (total time: 10 minutes).

For the post-RNA synthesis rinsing experiments, the RP_O_ was incubated with NTP in the reaction buffer for 10 sec before the reaction buffer was exchanged twice and finally replaced with imaging buffer, followed by the start of the acquisition. The buffer exchange procedure takes ∼40 sec to be completed before the start of the acquisition.

### Single-molecule data analysis

#### FRET pair localization and detection

The movies recorded on the camera were offline analyzed using the home-built Matlab routine Twotone-ALEX ^66^ to extract the intensities of co-localized donor and acceptor, i.e. FRET-pair. The following parameters from Twotone-ALEX were used to select only the FRET pairs formed by a single ATTO647N acceptor dye and a single Cy3b donor dye: channel filter as DexDem&&AexAem&&DexAem (colocalisation of the donor dye signal upon donor laser excitation, the acceptor dye signal upon acceptor laser excitation, and the acceptor dye signal upon donor laser excitation), a width limit between the donor and the acceptor between 1 and 2 pixels, a nearest neighbor limit of 6 pixels and a maximal ellipticity of 0.6 (ellipticity is defined as the ratio of the minor and the major axis of the ellipse). The traces extracted from the Twotone-ALEX analysis were then sorted to remove all the traces that displayed extensive blinking or multisteps photobleaching, i.e. that contain more than one donor or acceptor dye in the same diffraction limited intensity spot.

#### FRET efficiency characterization and Hidden Markov Modeling of the single-molecule FRET traces

The FRET efficiency dynamics for each FRET-pair was calculated with the standard formula 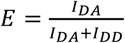, with *I*_*DA*_ and *I*_*DD*_ being respectively the intensity of the acceptor and of the donor upon donor excitation ^67^. The traces were analyzed through a modified version of the hidden Markov model (HMM) ebFRET software from Ref. ^68^, where only steps longer than 2 frames and separated from the subsequent step by more than twice the Allan deviation estimated at 5 frames were conserved ^69^ to be assembled into dwell time.

#### Characterization of the Δt_ITC6_, Δt_US_, Δt_PS_ and Δt_FS_ dwell time distributions

A detailed analysis of the dwell time distributions is provided in Ref. ^70-72^. Shortly, the distribution of τ are described by a probability distribution function with *m* exponentials:

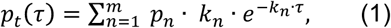

where *k*_*n*_ and *p*_*n*_ are the characteristic rate of the *m*^*th*^ exponential and its probability, respectively. The minimum number of exponential to fit the distributions was determined for each distribution by using the Bayes Schwarz Information Criterion (BIC) ^73^. We calculate the maximum likelihood estimate of the parameters (MLE) ^74^ by maximizing:

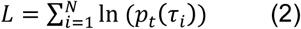

over the parameter set. Here the τ_*i*_ are the experimentally measured dwell times and *N* is the number of collected dwell times τ_*i*_. The error bars for each fitting parameters are one standard deviation extracted from 1000 bootstrap procedures ^75^. The ebFRET software ^68^ was also used to extract the peak positions of each FRET level, subsequently fitted with a Gaussian function, with the peak center and the standard deviation as free parameters (**Fig. S1D**, **Fig. S3A-D**).

